# Motion correction with subspace-based self-navigation for combined angiography, perfusion and structural imaging

**DOI:** 10.1101/2024.08.26.609650

**Authors:** Qijia Shen, Wenchuan Wu, Mark Chiew, Yang Ji, Joseph G. Woods, Thomas W. Okell

## Abstract

Motion artifacts are problematic in many MRI modalities. “Self-navigating” approaches are desirable, since no additional scan time or hardware is required. However, the generation of a navigator image, to estimate and correct motion, is difficult in cases where the tissue contrast is changing during the navigator acquisition window, such as in magnetization-prepared methods. Here we propose a subspace approach to reconstruct accurate navigators in the presence of time-varying tissue contrast and apply it to a combined angiography, perfusion and structural imaging method using a golden ratio 3D cones trajectory. This arterial spin labeling-based pulse sequence relies on subtraction of label and control images to isolate the relatively weak blood signal, making it particularly susceptible to motion corruption. An inversion pulse leads to time-varying tissue contrast across the readout train, but by reconstructing subspace coefficient maps directly, artifacts due to the varying contrast were alleviated. This resulted in high-quality navigator images that were subsequently registered to estimate and correct for motion. In addition, a split-update method was proposed to efficiently reconstruct from mismatched label/control k-space data with locally low rank regularization enforced on the difference image. The correction process was tested with numerical simulation and in vivo data from 8 healthy subjects with and without cued motion. In numerical simulation, the subspace-based navigator achieved an 84% reduction in RMSE of residual motion compared to without motion correction. In vivo, motion correction resulted in noise-like and background artifacts being greatly reduced and vessel sharpness being noticeably improved. Correlation of angiography, perfusion and structural images with motion-free reference images also increased by 159%, 53% and 12%, respectively, after motion correction. These results show that subspace-based navigators can effectively improve the motion robustness of MR imaging in contrast-varying acquisitions.

## 1 Introduction

MRI is susceptible to motion corruption due to prolonged scanning times. Subject motion, ranging from occasional nodding and coughing with healthy subjects to frequent and large-scale movements with disoriented patients, results in a decreased signal-to-noise ratio (SNR) and image artifacts. This degradation in image quality can negatively affect clinical diagnosis and increase costs due to the need for repeat scans (Andre et al., 2015; Godenschweger et al., 2016). Different MRI modalities are affected by motion to different degrees. Arterial spin labeling (ASL)-based blood flow imaging is particularly susceptible to motion corruption since it relies on the difference between two images in which the static tissue signal dominates. Therefore, any changes due to motion can lead to large artifacts that obscure the signal of interest.

A variety of MRI based techniques have been developed to estimate subject motion during scans. One stream of methods acquires dedicated navigator images during scanning to determine the motion status by comparing the navigators with a reference. Prospective motion correction (PROMO) (White et al., 2010; Zun et al., 2014), for instance, acquires 2D images on three orthogonal planes and compares them with the fixed reference navigator. Volumetric navigators were also used in some work (Frost et al., 2016; Grinstead et al., 2015; Shao et al., 2017), allowing a more straightforward comparison with the reference, while some methods perform motion estimation directly in k-space (Fu et al., 1995; van der Kouwe et al., 2006). However, insertion of additional navigators may alter the original contrast and increase scanning time. Self-navigated methods seek to correct for motion using the same data used for reconstructing the primary image. For example, joint motion estimation and image reconstruction was proposed for multishot structural MRI (Cordero-Grande et al., 2016). Navigators can also be reconstructed from the k-space center of radial data (Graedel et al., 2017).

However, the above mentioned methods could not be readily applied to correct for motion in a recently introduced ASL-based sequence called combined angiography, structural and perfusion radial imaging using ASL (CASPRIA) (Okell et al., 2022; Okell & Chiew, 2023), which has a global inversion pulse that leads to a time-varying signal during a train of spoiled gradient-echo readouts (Figure 1 and Figure 2a). Note that it shares similar form, a magnetization preparation followed by a long train of readouts, to other widely used sequences like MP-RAGE (Mugler & Brookeman, 1991) and fast spin echo (Mugler III, 2014). While the long train of readouts in each repeat potentially contains sufficient k-space information to reconstruct a single navigator, they cannot simply be combined because the image contrast varies across the navigator acquisition window due to the magnetization preparation. To address this, we propose to explicitly model the signal evolution across the readout train and incorporate it into a subspace-based reconstruction (similar to T2-shuffling (Tamir et al., 2017)) to reconstruct navigators with greatly reduced artifacts, thus improving registration accuracy.

**Figure 1.**
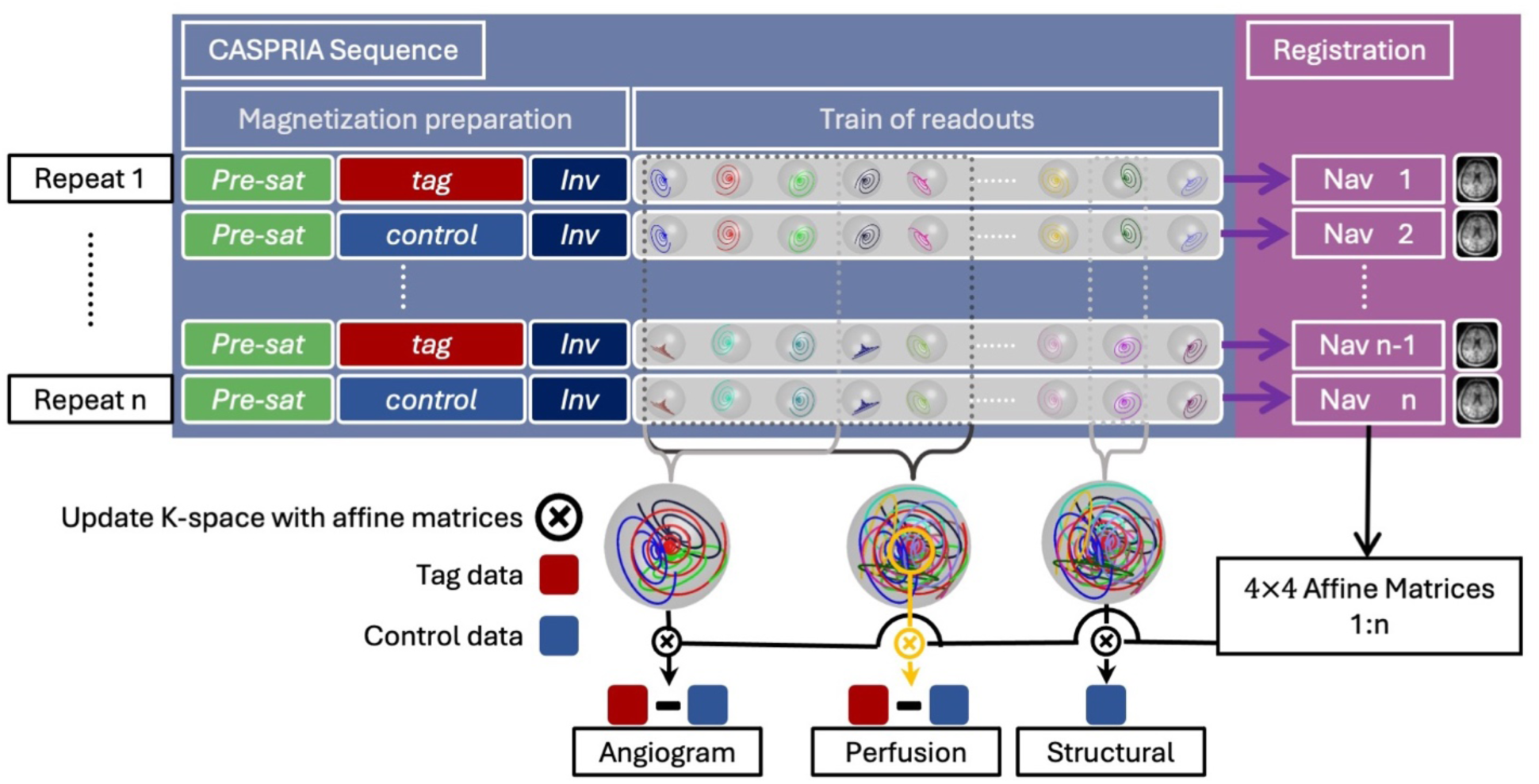
Overview of motion correction process using the CASPRIA sequence. Navigators 1 to n were reconstructed from the train of readouts in repeat 1 to n, respectively. Registration among all the navigators allows estimation of relative motion in the form of rigid transformation matrices. The matrices were then used to update the k-space trajectories and data, from which the motion corrected angiography, perfusion and structural images were reconstructed.

**Figure 2.**
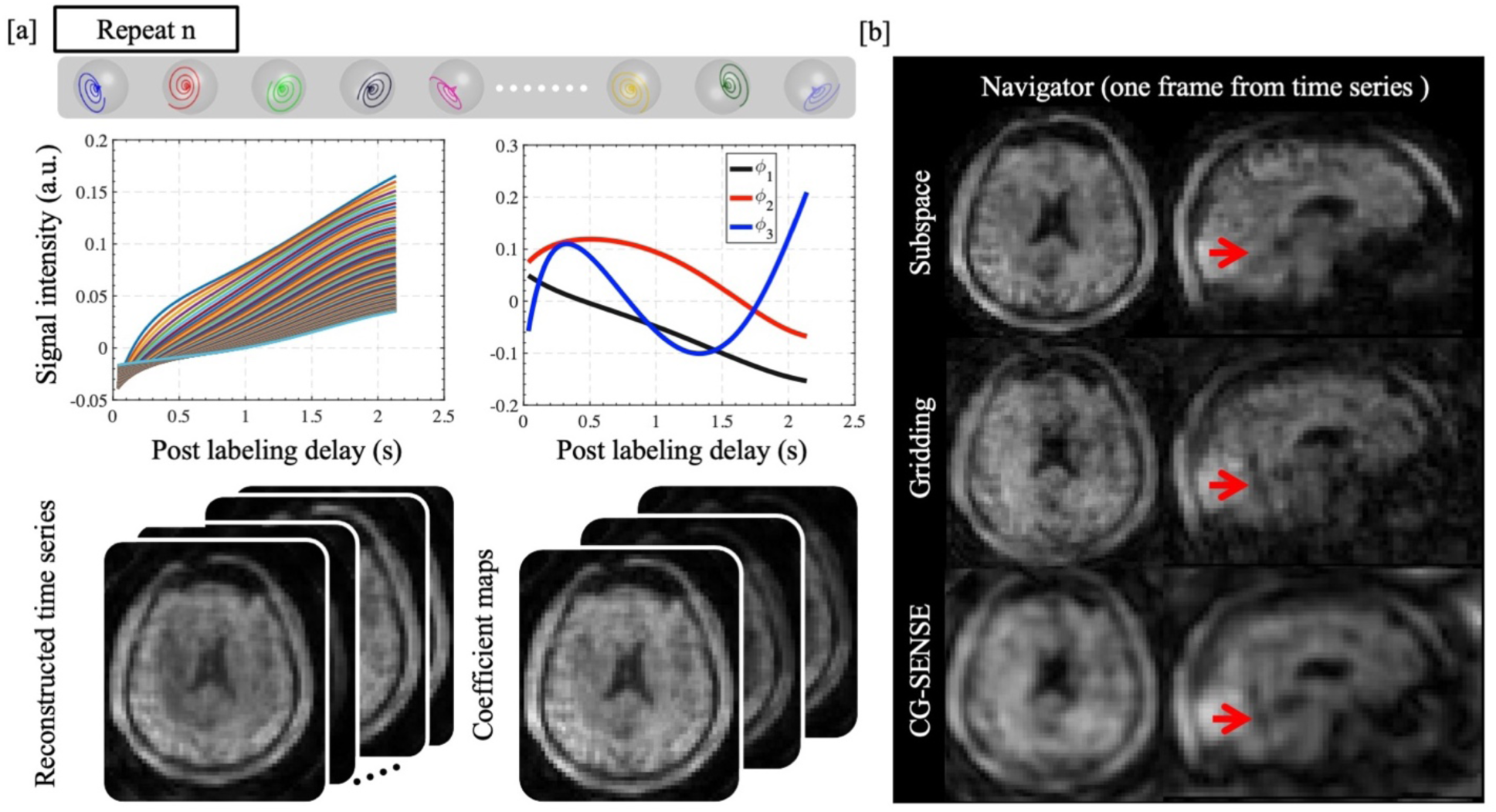
Reconstruction for subspace-based navigators. [a], for each repeat (without loss of generality, repeat n was shown as an example), the signal evolution of static tissue within each voxel follows a model dependent on sequence parameters and T1. Given the sequence, a dictionary of plausible 1D signal evolutions was simulated with various T1 values. The dictionary could be efficiently linearly compressed with 3 principal components (*ϕ*_1_ *ϕ*_2_ *ϕ*_3_). The **coefficient maps** α corresponding to the three principal components were reconstructed and then expanded to a full **time-series** of dynamic structural images using Eq. 2. Each frame of the time-series corresponds to the tissue contrast in one readout. [b], Comparison between an example navigator reconstructed with subspace approach, gridding and CG-SENSE methods. The frame corresponding to 960 *ms* after inversion in the **time-series** was selected as the actual navigator for motion correction. Artifacts pointed by red arrows could only be seen in gridding and CG-SENSE results and were speculated to be caused by contrast mixing.

Accurately estimated motion parameters using the subspace-based navigators were used to update k-space locations. However, methods like ASL require a subtraction between two sets of data with and without labeled blood (called label and control images), which is no longer feasible in k-space because the k-space sample locations are no longer matched after motion correction. Image space subtraction from separately reconstructed label and control images is possible, but it loses the advantage of artifact reduction by enforcing spatiotemporal regularization on the difference image. To mitigate artifacts in the subtracted image due to the different aliasing patterns of the strong background signal and residual motion, we propose a split-update method which jointly reconstructs the label and control images while imposing locally low rank (LLR) (J. Trzasko & Manduca, 2011) regularization on their difference.

The proposed subspace-navigator approach was applied to correct for rigid brain motion on the three modalities generated by the CASPRIA sequence: angiography, perfusion and structural images, while the split-update method was applied to the angiography and perfusion image reconstruction to mitigate subtraction artifacts. These methods were tested in numerical simulations and in a group of 8 healthy subjects, demonstrating the feasibility and significant improvement in image quality on all three modalities with the proposed motion correction approach.

## 2 Method

### 2.1 Pulse sequence design

As illustrated in Figure 1, the CASPRIA (Okell et al., 2022) sequence is an ASL-based sequence consisting of multiple repeats. Consistent with the general form of sequences our motion correction method can be applied to, each repeat is composed of a magnetization preparation followed by a train of readouts. The magnetization preparation part of the sequence consists of three sequential modules: water suppression enhanced through T_1_ effects (WET) (Golay et al., 2005) for pre-saturation of the imaging volume, pseudo-continuous arterial spin labelling (PCASL) (Dai et al., 2008) for labeling inflowing arterial blood, and a single adiabatic inversion pulse for background tissue suppression. Note that an interleaved scheme across repeats ensures the same k-space sample points are acquired for “label” and “control” images close together in time, which contain both static tissue and blood signal. Subtraction of label and control data isolates the labeled blood signal. The acquisition section is comprised of a train of readouts using non-Cartesian rotating cone trajectories (Shen et al., 2024) with 3D golden angle ordering (Chan et al., 2009). A number of readouts at adjacent acquisition timepoints are combined across all repeats to achieve sufficient k-space sampling for the reconstruction of dynamic images.

Assuming within-repeat motion is negligible, i.e., there is minimal motion during each readout train of around 2 seconds, a navigator was reconstructed from all the k-space data acquired during each repeat, as shown in Figure 1. The relative motion between repeats can then be estimated from registration among the navigators and is used to update the k-space trajectories and data. The motion corrected images (i.e., angiography, perfusion and structural images) were reconstructed from the updated k-space data.

As the inversion pulse, used for background suppression, introduces a continuously varying contrast during the train of readouts, grouping all readouts within a repeat and reconstructing with conventional methods like conjugate gradient (CG) SENSE (Pruessmann et al., 2001) or simple gridding (Fessler & Sutton, 2003) would generate images with significant artifacts and blurring (Figure 2b). Therefore, a subspace-based method was developed to accommodate for the signal evolution within each repeat.

### 2.2 Subspace-based reconstruction of navigators

Figure 2a illustrates the process of reconstructing navigator images using a subspace method for one repeat (without loss of generality, repeat n was used in Figure 2a). Firstly, we modeled the time-course of the tissue signal during the train of readouts after the inversion pulse. This time-course is approximately identical across repeat 1∼n (when ignoring the effects of the inflowing arterial PCASL labeled blood) and consists of a saturation-inversion T_1_ recovery curve, modulated by the readout excitation pulses.

Equation 1 describes how magnetization evolves from one readout to the next during one repeat. *i* denotes the *i*^*th*^ readout within the train of readouts and *α*_*i*_ denotes corresponding flip angle, which can vary across the readout train. Assuming equilibrium magnetization *M*_z_ = 1, *M_Zi_*^-^, *M_Zi_*^+^ represents the longitudinal magnetization immediately before and after *i*^th^ RF pulse respectively. *M_xyi_* represents transverse magnetization after *i*^th^ RF pulse. As TE is short in the sequence, T2* decay is not included in the model. TR represents the duration of single readout. Following Equation 1, the tissue signal, *M_xyi_* in one voxel can be sequentially calculated for each readout in a single repeat.

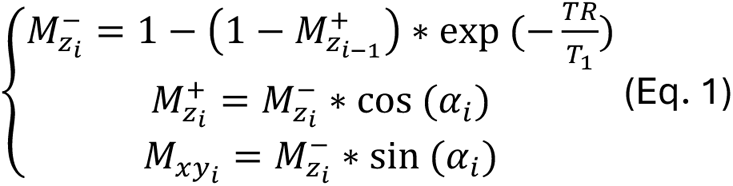

Given the sequence parameters TR and *α*_#_, the signal evolution is completely dependent on the T_1_ of each voxel. Therefore, a subspace was used to constrain the navigator reconstruction with this T_1_ dependent model. Firstly, a simulated dictionary of plausible signal time-courses was generated using Equation 1 with T_1_ ranging from 250ms -4000ms spaced by every 10ms. The parameters in Equation 1 followed the actual protocol described later. Secondly, principal component analysis (PCA) was used to linearly compress the dictionary. The first 3 principal components, Φ = [*φ*_1_ *φ*_2,_ *φ*_3_], represent 98.7% of the signal variance in the dictionary. The time-course of each voxel in the 4D images, *x*, can be represented as linear combination of the principal components, Φ, the weights of which are the coefficient maps, *α*, as proposed by Tamir et al., 2017:

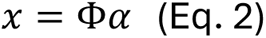

Given the subspace representation, the reconstruction of the coefficient maps from a single repeat can be formulated as:

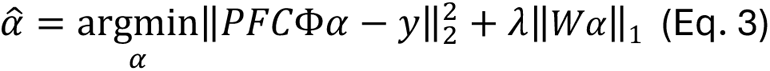

where P represents the k-space sampling, F is the non-uniform fast Fourier transform (NUFFT), C is the coil sensitivities, *W* denotes wavelet transform, and *λ* is the regularization weight.

After reconstruction, a 4D time series *x* was obtained for each repeat where each frame of *x* corresponds to the time of one readout after the single inversion pulse. The frame corresponding to 960 *ms* after inversion was selected as the actual navigator used for registration as it gives similar tissue contrast to MPRAEG (Mugler & Brookeman, 1991), which is known to be well suited to accurate registration.

Because of the correct modeling of the signal evolution across the readouts, the subspace reconstructed navigator demonstrated clearer anatomical structure and boundaries compared to the navigators reconstructed with gridding and CG-SENSE, as shown in Figure 2b.

### 2.3 Registration among navigators

The navigators reconstructed from each repeat were used to estimate subject movement. The linear registration tool FLIRT (Jenkinson et al., 2002) was used to register the navigator from each repeat to that of the first repeat as shown in Figure 1 and generated affine transformation matrix for each repeat. These affine transformation matrices were then used to update the k-space in each repeat. Specifically, an image space rotation in each affine matrix corresponds to a rotation of the k-space trajectory around the center, while an image space translation corresponds to a phase ramp in the k-space data. Each repeat was updated with the separately estimated rotations and phase ramps. The updated k-space was grouped for reconstruction of perfusion, angiography and structural images described below.

### 2.4 Split reconstruction of perfusion and angiographic images

Due to the high under-sampling factor used in CASPRIA acquisition, regularization is typically used to improve the conditioning of reconstruction. Conventionally, the label and control data are acquired with matched k-space trajectory and subtracted prior to the reconstruction of ASL perfusion and angiography images, i.e., only the difference image needs to be reconstructed. Therefore, a simple regularized reconstruction algorithm can be applied. However, with our proposed motion correction, the k-space trajectory in each repeat is updated with a different rotation matrix, so that the k-space trajectories in the label (odd repeat) and control (even repeat) acquisition are no longer matched, making direct k-space subtraction infeasible. Additionally, during testing we found that subtraction of separately reconstructed label and control images led to significant artifacts due to the different aliasing patterns of the strong static tissue signal; this prevented us from leveraging spatiotemporal correlations in the blood signal itself for regularization. Hence, we propose a new scheme where the label and control images are jointly reconstructed with a locally low rank (LLR) (J. Trzasko & Manduca, 2011) regularization enforced on their difference, as shown in Equation 4:

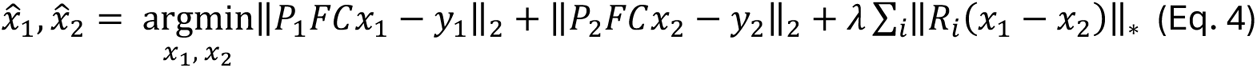

where *x*_1_, *x*_2_ are label and control images, respectively, *R_i_* is an operator for extraction of local patch *i*. and *P*_1_, *P_2_* represent sampling trajectories for label and control data. A split-update method adapted from POGM (Taylor et al., 2017) was used for optimization. To be more specific, the iterative optimization alternates between updating *x*_1_ and *x*_2_ as shown in Equation 5:

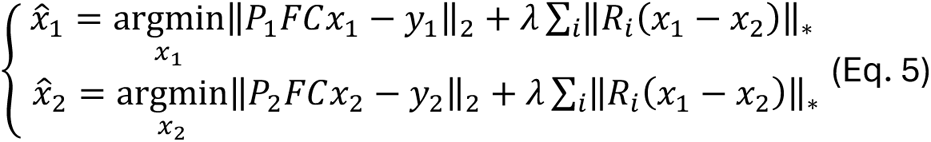

Where *x*_2_ and *x*_1_ are fixed when updating *x̑*_1_ and *x̑*_2_, respectively.

### 2.5 Subspace reconstruction for dynamic structural images

As shown in Figure 1, the dynamic structural image has the same temporal resolution as TR. The reconstruction still follows Equation 3 using the same subspace, except that *y* in Equation 3 now represents all data from PCASL control repeat, and the reconstruction target *α* has the spatial resolution same as angiography.

### 2.6 Experiments: Numerical simulation

To evaluate the motion estimation accuracy, numerical simulations were performed based on data from one scan with minimal subject movement. The acquisition protocol used in simulation was the same as the in vivo scanning protocol described below. A motion trajectory specifying rotation and translation parameters for each repeat, shown as the black curves in Figure 3a (the complete set of parameters can be found in Supplementary Figure S1), were used to corrupt the k-space data, simulating head movements during the scan. The navigator image for each repeat was reconstructed and registered following the methods described above. The residual uncorrected motion was calculated by multiplication of the estimated affine matrix and the ground-truth motion matrix, such that perfect motion estimation would result in an identity matrix. Residual motion parameters (rotation and translation on 3 axes) were then calculated from the residual affine matrix using the ‘avscale’ function in FLIRT (Jenkinson et al., 2002). For comparison, navigator images were also reconstructed with gridding (Fessler & Sutton, 2003) and CG-SENSE (Pruessmann et al., 2001) and evaluated in the same manner to help assess the potential benefits of the proposed subspace approach.

**Figure 3.**
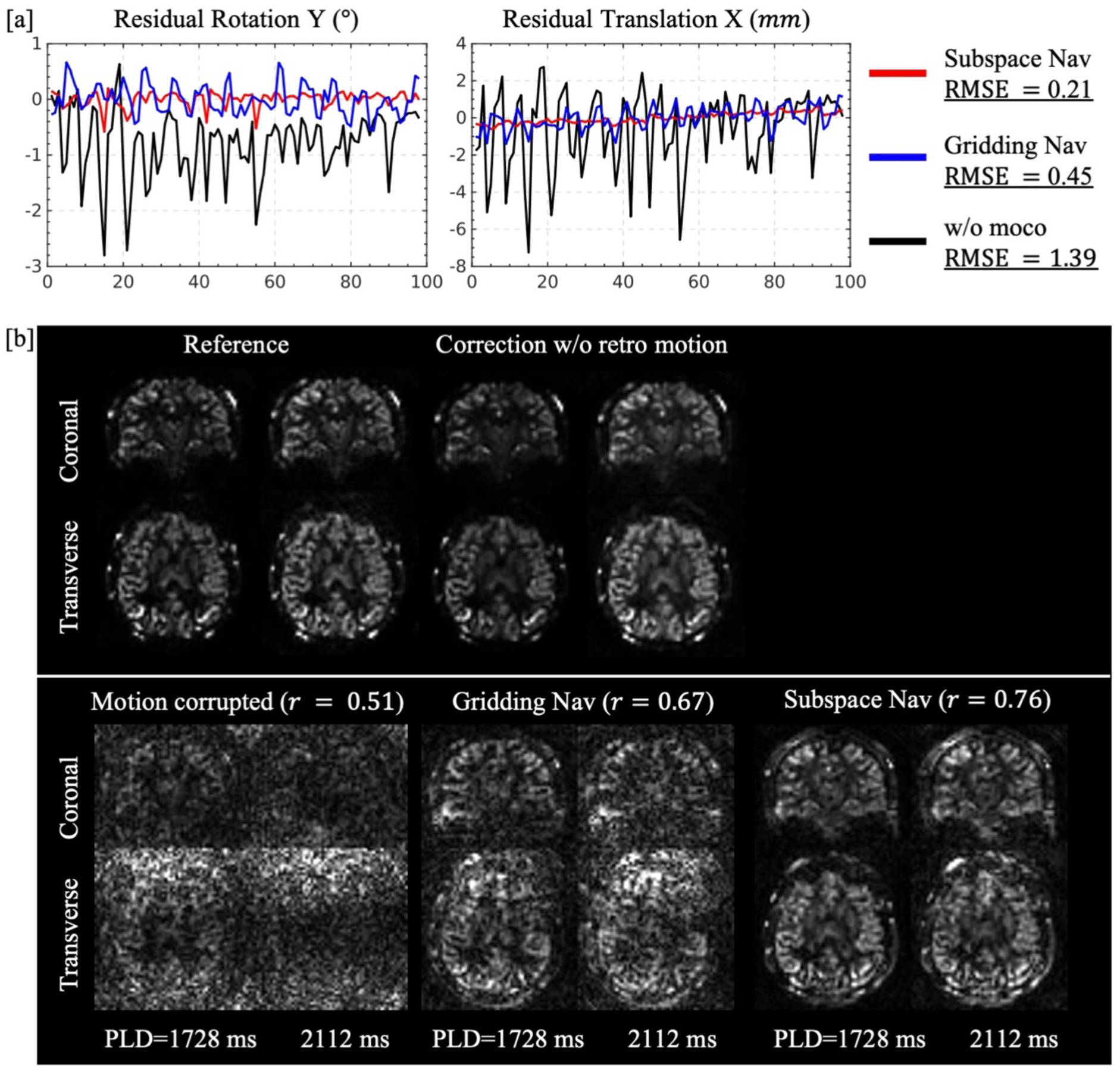
Numerical simulation with motion parameters retrospectively added onto a dataset with minimal subject motion. [a], Motion parameters retrospectively added onto the motion-free k-space data (black curve), and residual motion estimated with navigators reconstructed by gridding (blue) and subspace (red) methods. RMSE of 6 parameters of the residual motion was calculated for each case. [b], Perfusion images calculated from data with retrospectively added motion. Each dynamic perfusion image has six temporal frames, the last two of which are shown. The motion correction pipeline with subspace derived navigators was applied to the data without retrospectively added motion first to test robustness. Correlation (*r*) was calculated between the two frames of the reference image and those corrupted by motion, corrected with gridding navigators and subspace navigators.

Since perfusion images are most sensitive to motion, due to the weak blood signal relative to static tissue, simulated perfusion images were reconstructed from k-space data without motion, with added motion and with correction on added motion to qualitatively assess the efficacy of the proposed method.

As both the rotation and translation have similar scales, root-mean-square error (RMSE) of all residual rotation and translation parameters from all repeats were calculated to quantitatively evaluate the motion estimation accuracy.

### 2.7 Experiments: In-vivo scanning protocol

#### 2.7.1 Acquisition parameters

To assess the proposed motion correction method, a total number of 8 subjects were scanned on a 3T Prisma scanner (Siemens Healthineers, Erlangen, Germany) using a 32-channel receive-only head coil under a technical development protocol agreed by local ethics and institutional committees. Two back-to-back scans with the same protocol were performed on each subject. In the first scan, subjects were instructed to move their head to follow a marker on the screen, which moves every 20 s (details can be found in Supplementary Figure S2). In the second scan, subjects were instructed to stay still. We termed the data for the first scan as “cued motion data”, and the second as “reference data”.

For each scan, the protocol followed the settings used in our previous study (Shen et al., 2024): briefly, after each PCASL preparation and single inversion pulse, a train of 144 readouts was acquired within each repeat. A quadratically varying flip angle scheme was used in each readout train, ranging between 3°-12°. The temporal resolution used to reconstruct perfusion and angiographic images was 384 ms and 192 ms, respectively.

Dynamic structural images were reconstructed using the subspace approach with a temporal resolution equal to the readout TR (16 ms). 12 frames, 6 frames and 144 frames were reconstructed for angiography, perfusion and dynamic structural images respectively.

An additional MPRAGE (Mugler & Brookeman, 1991) image was acquired for each subject as reference for evaluating motion correction accuracy. The sequence used TI = 904 *ms*, TE = 3.4 *ms*, echo spacing = 8.8 *ms*, turbo factor = 128, flip angle = 8°,and spatial resolution = 1.7 mm isotropic.

#### 2.7.2 Reconstruction parameters

Coil compression (Buehrer et al., 2007) was used to reduce the original 32 channel data to 8 channels for reconstruction. Coil sensitivity maps were estimated from a pseudo-structural image (reconstructed by regridding averaged label and control data from the final 360 ms of the readout train where the static tissue signal is more consistent) using the adaptive combine method (Walsh et al., 2000) prior to motion correction. Despite existed motion, we found that coil sensitivity maps can still be reliably estimated from the pre-correction pseudo-structural image. For both navigator and dynamic structural image reconstructions, 3 principal components were used and *λ* = 1*e* − 4 was empirically chosen as the weight for wavelet regularization. For perfusion and angiography reconstructions, *λ* = 7*e* − 2 was chosen empirically as the weight for LLR regularization. The locally low rank patch size was 5 × 5 × 5 voxels.

The reconstruction matrix sizes for both angiography and structural images are 186 × 196 × 150 with isotropic spatial resolution 1.13 *mm*. The navigator and perfusion images share same matrix size 62 × 66 × 50 and isotropic spatial resolution 3.39 *mm*.

### 2.8 Quantitative evaluation

To investigate the effectiveness of our motion correction method quantitatively, Pearson correlation coefficient was calculated between images with and without cued motion before and after applying motion correction, where the reference images are the data without cued motion and reconstructed without the motion correction pipeline. To align the position of the two datasets without introducing blurriness due to image space interpolation, a pseudo-structural image was reconstructed for data acquired with and without cued motion. A transformation matrix was extracted by registering the pseudo-structural image of motion corrected data to reference data. The same transformation matrix was then multiplied with the affine matrices generated by the registration step for updating the k-space before reconstructing the final angiography, structural and perfusion images, such that the motion corrected images were aligned with the reference images.

A brain mask, used to calculate correlation for structural and perfusion images, was extracted using BET (Smith, 2002), while a dilated vessel mask, used to assess the angiographic data, was extracted following the steps described in the original 4D CAPRIA paper (Okell & Chiew, 2023). The correlation between the cued motion data and reference data was then calculated within the relevant mask for each frame separately. In addition, the dynamic angiogram within the vessel mask was also fitted with FABBER (Chappell et al., 2009) and the resulted transit time was displayed to demonstrate the effectiveness of motion correction.

Multi-way ANOVA test was performed on the fisher transformed correlation coefficients to assess the statistical significance of differences before and after gridding and subspace-based motion correction, with subject number and post labeling delay (PLD) as additional factors.

## 3 Results

### 3.1 Numerical simulation

Results from the numerical simulations are shown in Figure 3. The retrospectively added motion trajectory is plotted together with the residual motion after correction by navigators reconstructing using either the subspace or the gridding method. Motion could be largely estimated by registering the gridding reconstructed navigators, reflected by the 67% reduction of RMSE of motion parameters compared to without motion correction. However, mixture of contrast and unclear brain/tissue boundaries, as shown in Figure 2b, limited the motion estimation accuracy using gridding navigators. The subspace-based reconstruction, however, greatly reduced artifacts in the navigator images by correctly modeling the signal evolutions across time. Additional LLR regularization further reduced the non-Cartesian trajectory induced artifacts and improved the boundary clarity. Therefore, the retrospectively added rotation and translation were accurately estimated by registering the navigators reconstructed by the subspace method, as depicted by the red curve in Figure 3a, which achieved an additional 53% reduction in RMSE of residual motion compared to the gridding reconstructed navigators.

The visual quality of the perfusion images, especially last two frames, serves as a good indicator of how good a navigator was at accurately estimating motion because perfusion images are particularly susceptible to motion corruption and the last two frames have the lowest perfusion-to-tissue relative signal intensity ratio. Before adding retrospective motion to the raw data, our motion correction pipeline was first tested on the original data to ensure data quality was not degraded by the correction in cases with minimal motion, as shown in Figure 3b “correction w/o retro motion”. The high similarity between the “reference” and “correction w/o retro motion” demonstrates the stability of the proposed method. Simulation with retrospective motion addition was then performed. As shown in Figure 3b, the motion corrupted perfusion images were dominated by artifacts, almost without discernable brain structure. After correction with motion estimated from the gridded navigators, more detail was recovered. The tissue structure, however, was still mixed with artifacts. In comparison, the perfusion images corrected with subspace reconstructed navigators revealed greatly improved tissue detail with the fewest artefacts.

The improved correlation (*r* = 0.76 in Figure 3b) between motion corrected and motion free reference perfusion images also demonstrated that subspace-based navigators could correct simulated motion more accurately than gridding-based navigators (*r* = 0.67).

### 3.2 In-vivo perfusion results

Consistent with numerical simulation, the subspace-based navigator can also accurately estimate subjects’ movement on in-vivo data with cued motion. As shown in Figure 4a, a subject moved intentionally during the scan, resulting in significant artifacts covering the whole brain. The tissue structures were mostly recovered after motion correction. Figure 4b showed another subject with more severe motion corruption. The data was virtually overwhelmed by motion induced artifacts before correction, preventing any kind of qualitative assessment. Our correction method successfully recovered tissue perfusion information. In the motion corrected images, more artifacts are observed at PLD=1728 ms compared to 1344 ms likely due to ineffective suppression of background tissue signals because of T1 relaxation (see Figure 2a).

**Figure 4.**
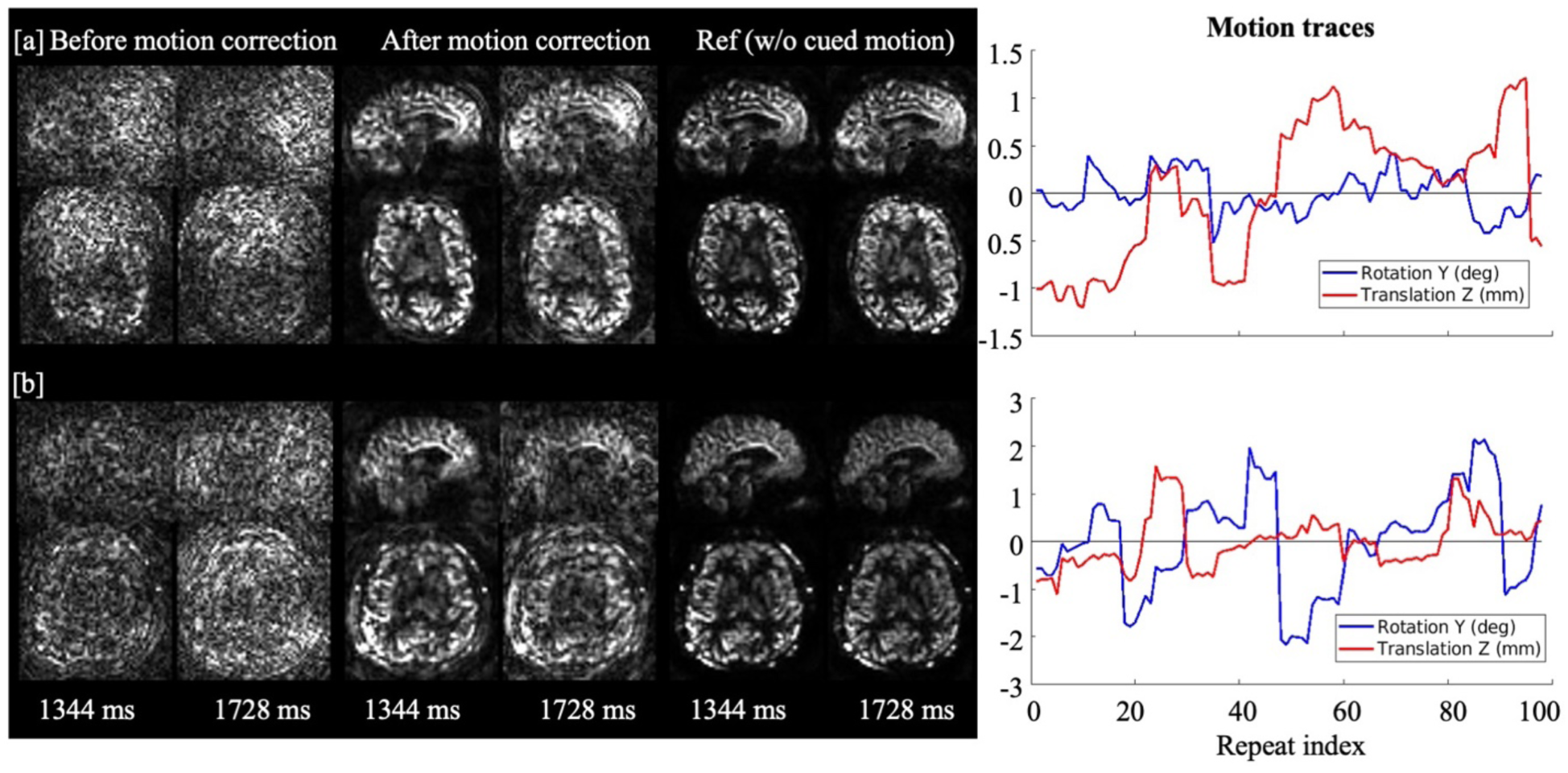
Perfusion images from the data acquired with cued motion compared to reference data. [a], this subject moved during perfusion scan with moderate amplitude and frequency as plotted on the right. The quality of the perfusion images improved after motion correction. [b], Data acquired from a different subject with more severe motion. Artifacts in the perfusion images were greatly reduced as well after motion correction, especially at PLD=1728 ms.

### 3.3 In-vivo angiography results

Despite its strong signal intensity compared to perfusion imaging, angiography is also affected by motion because visualizing thin vessels with fidelity requires accurate alignment between repeats. The images before motion correction in Figure 5a demonstrate an example of the effects of motion on an angiogram acquired with cued motion. The thin vessels in Figure 5a were blurred and mixed with background tissue signal due to motion at PLD=480 ms. As the background tissue signal recovers with advancing PLD, the vessels become obscured by motion artifacts. As exemplified by the enlarged green box in Figure 5a, almost no discernible vessels could be identified before correction, but they were largely restored after correction.

**Figure 5.**
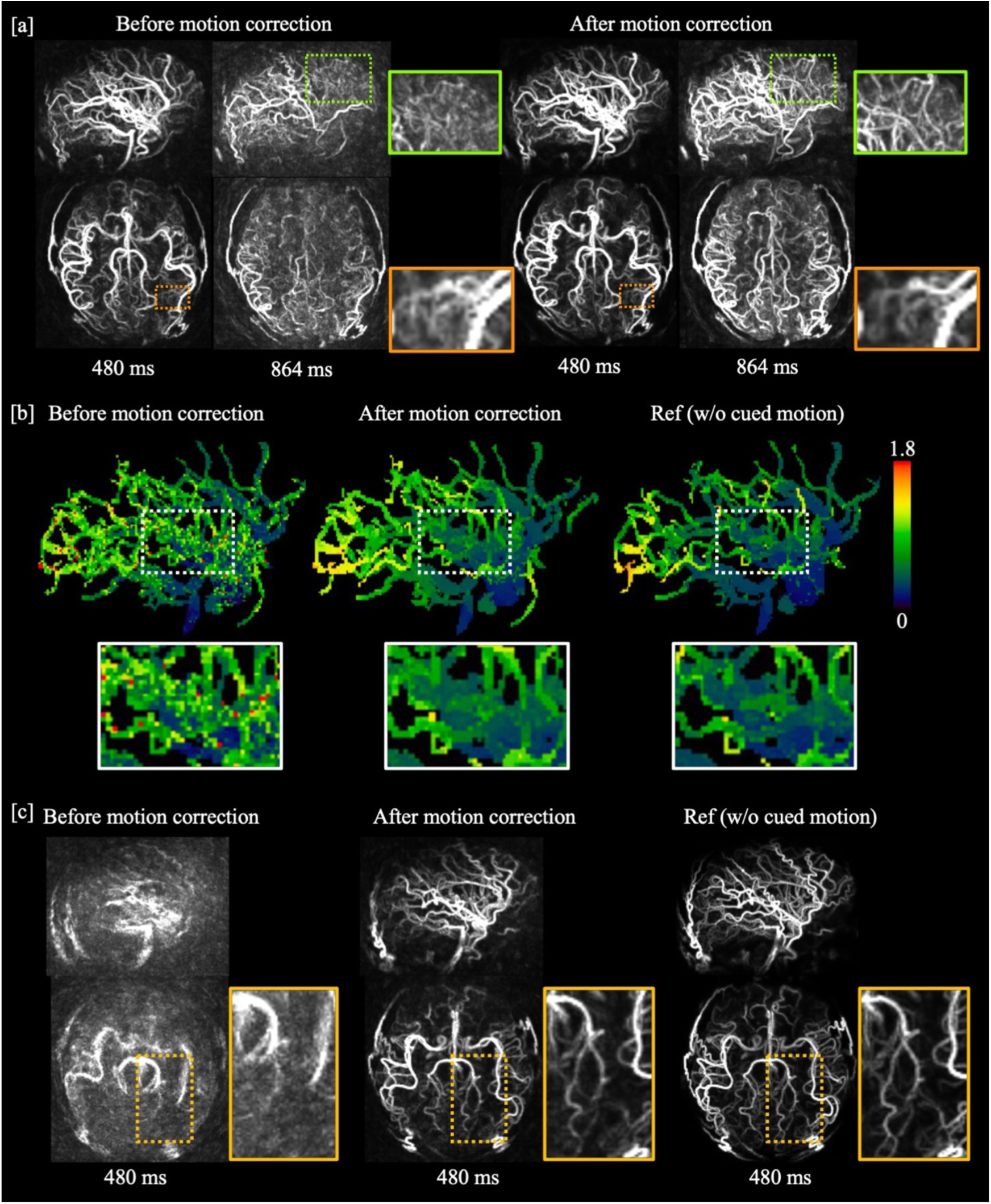
Comparison of angiography images before and after motion correction. [a], Angiography images still preserved fidelity under moderate motion. The effect of motion was mainly manifested as blurring of thin vessels at short PLDs when the background tissue signal was well suppressed. As the background suppression gets weaker due to recovery of the static tissue signal (e.g. at PLD=864 ms), motion resulted in a complete loss of visible small vessel signal, but this was recovered with the proposed motion correction approach. [b], Comparison of transit time estimation from the angiogram in [a] before and after motion correction. Spurious estimation caused by motion was mostly removed in motion corrected data. [c], A different dataset showing an angiogram affected by very large motion, which was still well recovered after correction. The corresponding motion traces can be found in Supplementary Figure S3.

The misalignment between repeats not only caused impairment of vessel visibility, but also degradation of quantification results. As shown in the white box in Figure 5b, the transit time estimated from motion corrupted data contained many more spurious values than the motion corrected one and deviated from the reference result.

During more extreme motion in a different subject, as displayed in Figure 5c, this angiogram suffered from almost complete loss of visible vessels. The motion traces for this severe case were plotted in Supplementary Figure S3. However, fine details were recovered in the motion corrected images, further confirming the motion estimation accuracy of the proposed method.

### 3.4 In-vivo structural results

The structural images were reconstructed using the subspace method, and the frame corresponding to 960 ms after inversion pulse was compared to MP-RAGE in Figure 6. As shown in shown in Figure 6a, moderate motion resulted in relatively minor blurring of tissue structures which were visibly improved using the proposed motion correction method. In Figure 6b, the data affected by strong motion resulted in much greater blurring. Nevertheless, motion correction greatly improved the image quality, and the tissue structure details were recovered.

**Figure 6.**
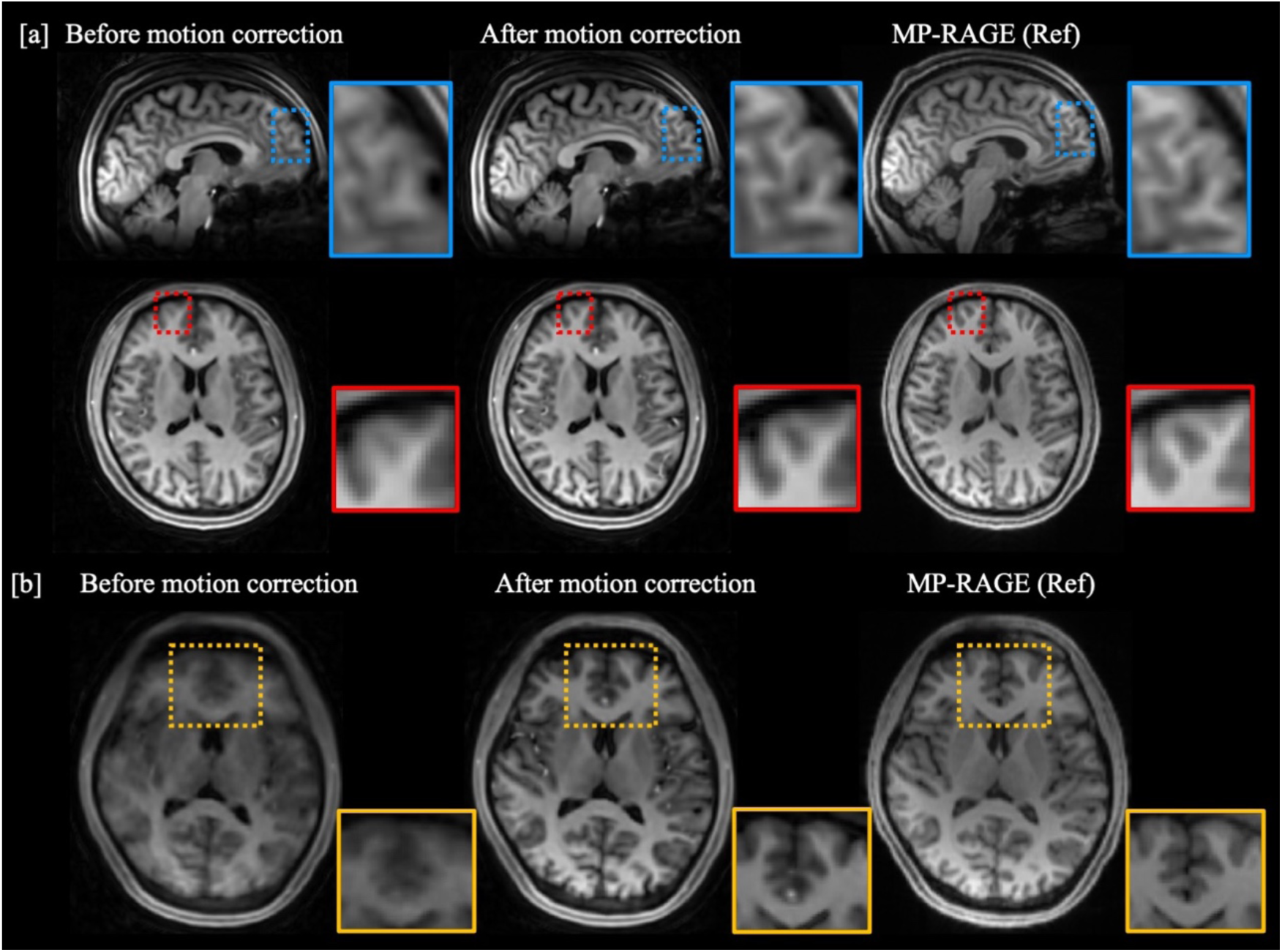
Comparison of structural images before and after motion correction. The data was acquired with cued motion. [a], Moderate motion caused blurriness in brain tissue structures. The structural sharpness was noticeably improved after motion correction, as shown in the blue and red boxes. [b], Structural images affected by strong motion lost much more structural detail but could still be corrected with high quality compared to the reference motion-free MP-RAGE image, as shown in the yellow box.

### 3.5 Robustness to mild motion

One important motivation for this study originated from an observation of previous experiments that, even if a subject intends to stay still, mild and unintentional motion could still degrade the quality of perfusion images. In Figure 7, we demonstrated two such cases to prove that our subspace-based correction method is also robust when motion is small. These two cases were noticed being corrupted by motion artifacts among the data acquired without cued motion (reference data session). For one subject, the head position drifted during scanning accompanied by sudden rotations. The motion induced artifacts were mainly concentrated in frontal regions and exhibited similarity to noise. Motion induced artifacts for the second subject were more obvious. In both cases, the perfusion images were greatly improved after motion-correction, so they were still included as reference images when calculating the correlation with the case with cued motion in Figure 9.

**Figure 7.**
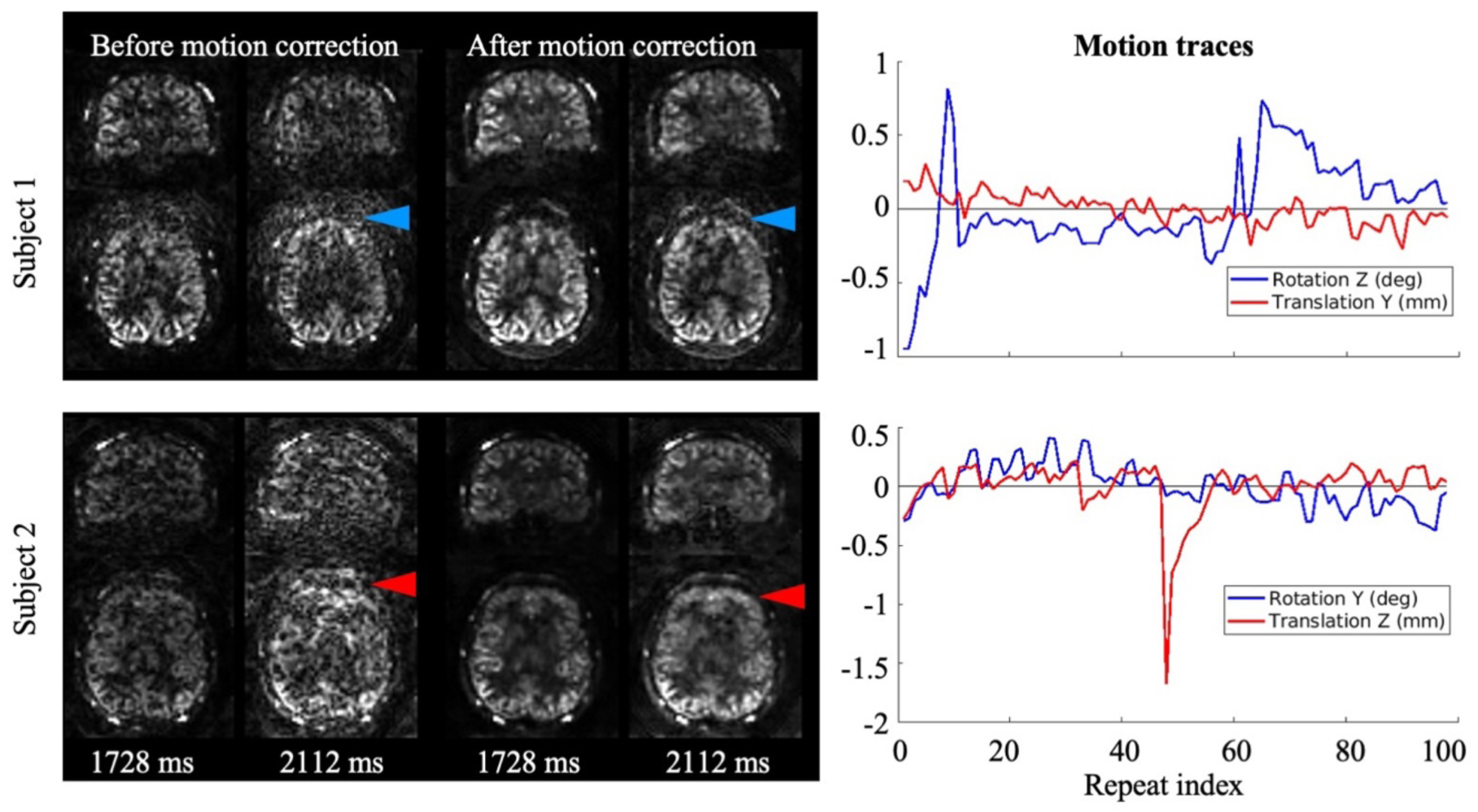
Perfusion images before and after motion correction for two datasets acquired without cued motion, but which still demonstrated some motion corruption. The images after correction showed noticeable reduction of artifacts in the frontal area (blue and red arrows).

### 3.6 Qualitative comparison between gridding and subspace-based correction

Image quality of in-vivo results corrected by subspace-based navigator also notably surpassed those by gridding-based navigator. Following numerical simulation, we also used perfusion results at long PLD as the indicator due to their high sensitivity to k-space mismatch. In Figure 8, we compare the reconstruction results based on motion estimated by subspace navigators with that by gridding navigators using cued-motion data from one subject. Without motion correction, the perfusion images are dominated by artifacts without any discernible structures. Although many tissue structures were recovered after correction with the gridding-based navigators, the image quality was inferior to that of the subspace-based correction. This result is consistent with numerical simulation and confirms that subspace-based navigators can provide more accurate motion estimation than gridding-based navigators. The quantitative comparison between results corrected by two navigators were also performed on the group level and will be displayed in the next section.

**Figure 8.**
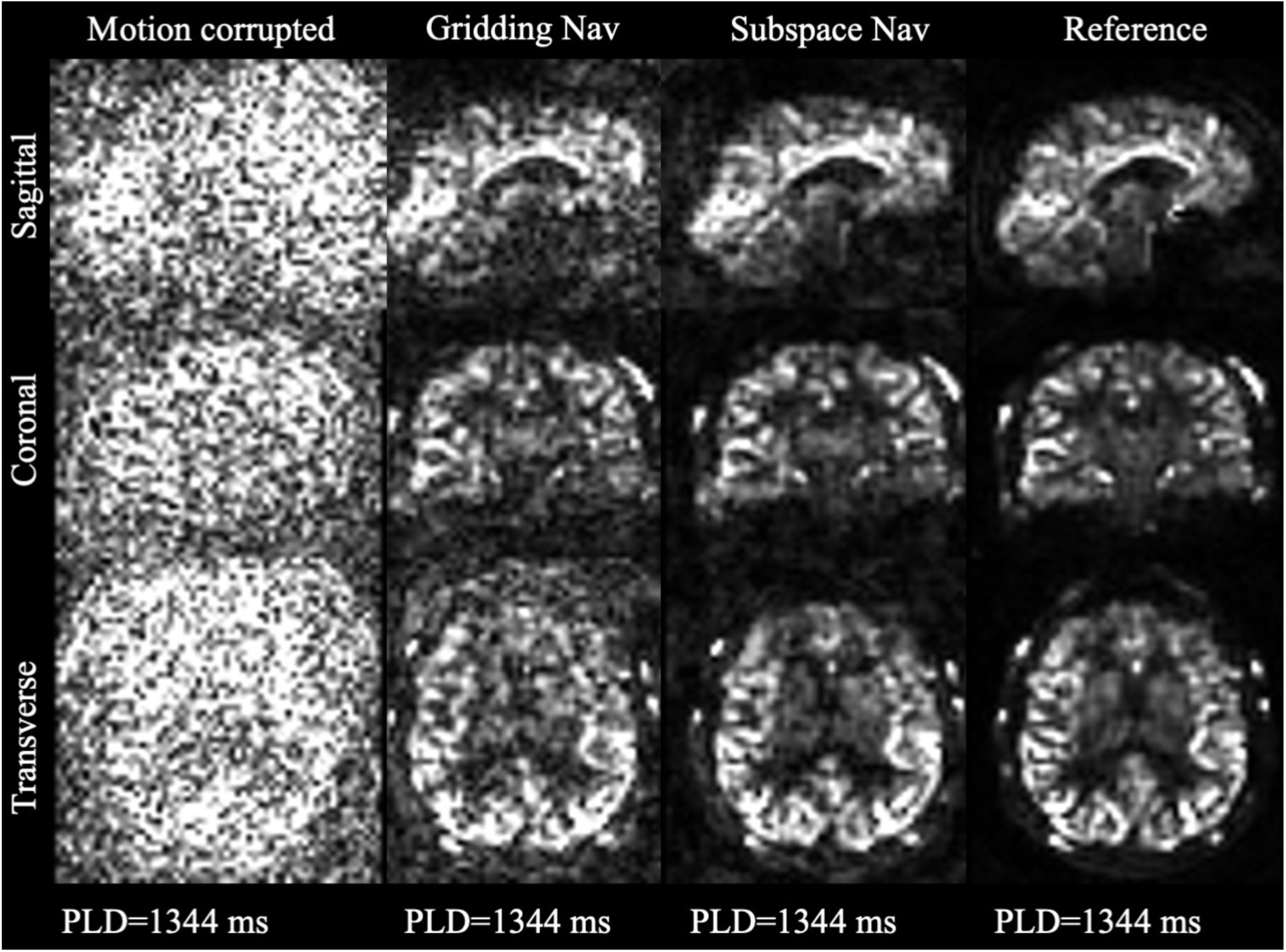
Comparison of one frame (*PLD* = 1344 *ms*) of the perfusion images from the cued motion scan with gridding-based and subspace-based navigators. From left to right are perfusion images reconstructed without motion correction, motion-corrected with gridding navigator, motion-corrected with subspace-based navigator, and motion-free reference data in a single subject.

### 3.7 Quantitative results

Correlation coefficients calculated within the group of subjects on all three modalities are presented in Figure 9. Figure 9a shows the correlation of the motion corrected structural images with reference. As comparison on all 144 frames of dynamic structural images is computationally demanding, a representative frame corresponding to 960 ms after inversion pulse with rich tissue contrast is chosen for evaluation. Consistent with the qualitative results, the structural images are less sensitive to motion compared to the perfusion and angiography images and thus have higher correlation to reference data. As subjects moved differently in the scanner, the correlation without motion correction has large variance. After gridding-based motion correction, the calculated correlation to reference has a higher average with limited variance, which was further improved after using subspace navigators.

**Figure 9.**
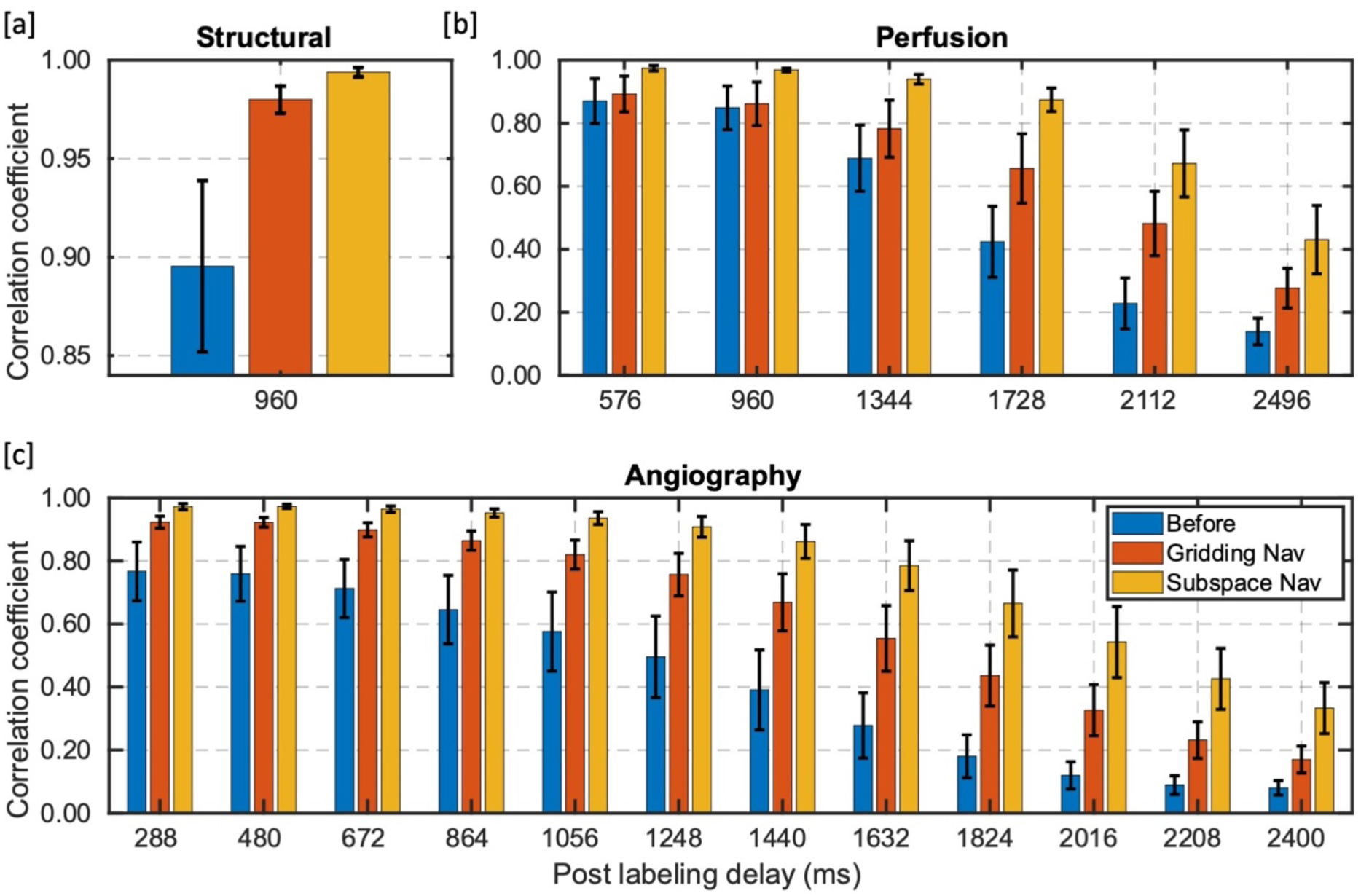
Comparison of correlation coefficient before and after gridding-based, subspace-based motion correction. Mean and standard deviation of correlation on the group of 8 healthy subjects were plotted per frame. Note that the mean values were plotted as columns where the standard deviation in the form of error bars were overlaid. [a], Bar plot of the correlation of structural images. The correlation was calculated on the frame with PLD = 960 *ms*. Subspace-based navigator improved 12% and 1.4% of correlation compared to before and after gridding-based correction respectively. [b], Bar plot of the correlation on all 6 frames of the perfusion. There were 53% and 16% improvement of correlation on average using subspace-based motion correction when compared to before and after gridding-based correction. [c], Bar plot of the correlation on all 12 frames of the angiographic images. 159% and 11.2% improvement of correlation were achieved on average with subspace-based correction versus before and after gridding-based correction respectively across all subjects and frames.

Correlation coefficients of the perfusion images is shown in Figure 9b, where r-value of all 6 reconstructed frames is presented. The overall correlation coefficients to reference data were lower than the structural data as perfusion signal is weaker and can easily be corrupted by static tissue when label and control data is mismatched. Nevertheless, the subspace-based correction still led to 53% increase in correlation to reference compared to without correction across all frames and subjects, and 16% additional improvement compared to gridding-based navigators. As was observed in the CAPRIA paper (Okell & Chiew, 2023), the standard deviation of correlation was increased in the last two frames but not necessarily due to uncorrected motion. The decreased correlation can also be attributed to weaker background suppression at long PLD, smaller perfusion signal and more significant effects of noise as well as residual static tissue artefacts. In Figure 9c, the angiography results also achieved significant improvements in correlation to reference, 159% and 11.2% compared to without correction and with gridding-based correction respectively. Overall, a statistically significant (*p* < 0.001) increase in correlation to reference data was found in subspace-based correction compared to gridding-based correction and, naturally, without motion correction.

## 4 Discussion

Here, we proposed a method for reconstructing robust navigators for MRI acquisitions in which the contrast is changing over time. We showed that a subspace approach reconstructs navigators with better quality than conventional gridding or CG-SENSE methods when the contrast changes and gives much more robust motion estimation. We further proposed that, for methods which rely on subtraction, a split-update reconstruction can be used to incorporate spatiotemporal regularization on the difference signal whilst accounting for the mismatched k-space sample locations between label and control conditions (e.g., after being updated with different motion parameters). We demonstrate these improvements on the CASPRIA sequence in which structural, angiographic and perfusion images can be obtained from a single scan, showing large reductions in motion artifacts and significant improvements in correlation to motion-free reference.

This work extended the previous development of the CASPRIA sequence (Okell et al., 2022; Okell & Chiew, 2023) and the optimized cone trajectory (Shen et al., 2024), with a focus on improving the motion robustness with self-navigation. The self-navigated motion correction approach possesses several advantages: 1) the method does not prolong the total scanning time, which is crucial in clinical settings. Adding a navigator for each repeat would add a considerable amount of scan time considering the total number of repeats reaches ∼100; 2) the original contrast is unaffected, avoiding the possibility of introducing unexpected signal change or artifacts; 3) the proposed motion correction method also only leads to a modest increase in reconstruction time and can be inserted as an initial step to existing reconstruction pipelines. Reconstruction and registration for each navigator using the subspace-based method takes 10-20 mins, and these could be run in parallel for each repeat. The memory and time consumption for the split-update reconstruction for each modality increased by less than twice. In contrast, the joint parameter estimation and reconstruction method (Cordero-Grande et al., 2016) would be computationally infeasible when dealing with high-resolution 4D dynamic images with ∼100 sets of motion parameters.

The techniques developed, however, are not confined to the CASPRIA sequence itself and are suitable for translating to different applications. The subspace-based navigator could see its application in cases with long trains of readouts and varying contrast. For example, MP-RAGE (Mugler & Brookeman, 1991), the signal curve of which is plotted in Supplementary Figure S4, MPnRAGE (Kecskemeti et al., 2016)) and fast spin echo sequences (Mugler III, 2014; Tamir et al., 2017) could benefit from subspace navigators, as incorporating the signal model compensates for undersampling common in navigator reconstructions and correctly matches signal and contrasts at different timepoints, which, in turn, helps reduce artifacts in reconstructed navigators and improves the registration accuracy. Nevertheless, it is not trivial to incorporate the proposed subspace-based navigator into existing sequences for motion correction as the spatial encodings might need to be reordered to have sufficient sampling after each magnetization preparation phase.

The proposed split-update method is also generalizable for reconstruction of the ASL signal in other sequences. Previous ASL methods subtract the label and control signal either in k-space prior to reconstruction (J. D. Trzasko et al., 2011) or in image space post reconstruction (Günther et al., 2005). K-space subtraction simplifies reconstruction by optimizing for the difference image only but is incompatible with any mismatch in k-space sampling points. Image space subtraction affords more flexibility in k-space trajectory but loses the benefit of image space regularization and reduces SNR. Our split-update method combines the advantages of the two. Integrating the two variable updates into a gradient method (e.g., POGM) resulted in convergence using iterations similar to single variable updates. Although image space subtraction methods have also been proposed by others (Rapacchi et al., 2014; Yu et al., 2012), they reconstruct each frame separately. Our method reconstructs the whole timeseries and incorporates LLR constraints (J. Trzasko & Manduca, 2011) for effective denoising across frames. We also considered using the ADMM optimization algorithm (Boyd et al., 2011) for joint label and control reconstruction, the formulation of which is in the Supplementary Reconstruction section. But in preliminary experiments we found it difficult to adjust the parameters for results comparable to the split-update method. The data consistency step within each iteration is also iterative which makes reconstruction time for high resolution dynamic angiograms prohibitively long (several days).

To reduce computational time and memory consumption, the sensitivity maps were not transformed accordingly within each repeat when incorporating the motion parameters into the reconstruction for the in-vivo results. To be more specific, the computational time and memory consumption scales linearly with the number of repeats (i.e., ∼100) when incorporating the transformed sensitivity maps for each repeat, and therefore, prohibits running reconstructions on dynamic perfusion or angiography images. This approximation, however, did not seem to severely impact the quality of the motion corrected images according to our results, as sensitivity maps are generally smoothly varying. This approximation still appeared to hold even with large head movements. For example, in Figure 5c, the angiogram after motion correction showed high fidelity and sharpness compared to the reference while the rotation reached 15°.

As the complete train of readouts within each repeat was used to reconstruct one navigator, the minimal interval our method could discern between motion states was ∼2s. Nevertheless, a report examining fMRI data of 42874 subjects in UK Biobank (Hess, 2022) showed that the vast majority moved slower than 0.5 deg/s in rotation and 0.5 mm/s. Therefore, we believe the 2s interval used in this work could already correct for a considerable amount of motion.

In addition, within-repeat motion estimation might also be possible with the help of our subspace reconstruction method. Similar to the cloverleaf (van der Kouwe et al., 2006) method, subspace navigators could be used as a reference for each repeat while the motion status of each individual readout could be estimated by comparing to the reference in k-space, and achieve more sophisticated motion correction.

Another source of imperfect correction originated from the insufficient background suppression towards the end of the readout train as static tissue recovered after the inversion pulse. Insertion of an additional inversion pulses during the train of readouts would help to reduce the intensity of static tissue at long PLDs. Subspace-based navigators could still be reconstructed by explicitly modeling the additional inversion pulses. In conventional self-navigated ASL motion correction methods, background suppression is deliberately suboptimal for higher tissue contrast in the navigators (Zun et al., 2014). However, this situation can be avoided, because the subspace-based method reconstructs the whole time-series and the frame with best tissue contrast can be selected as the navigator.

As a preliminary study, a group of 8 healthy subjects were scanned with and without cued motion. A broader comparison within a large group should be carried out in the future, examining our method in correcting for involuntary motion of healthy subjects as well as patients with more severe head motion.

## 5 Conclusion

In this work, we proposed a subspace-based method for reconstructing navigators in acquisitions with changing contrast and showed that the reconstructed navigators have better quality and registration accuracy than gridding with contrast-mixed signal. A split-update algorithm was also introduced to reconstruct from mismatched k-space sample locations while applying LLR regularization on the difference image. Both methods were applied to the CASPRIA sequence for motion corrected angiography, structural and perfusion images. The motion correction pipeline was evaluated comprehensively with numerical simulation and achieved 84% reduction in RMSE of the residual motion parameters. In scanning of healthy subjects, it also noticeably reduced motion artifacts. Compared to the reference, correlation of the perfusion, structural and angiography images between the motion corrected and reference data was increased by 53%, 12% and 159%, respectively, after motion correction.

## Data and code availability

The code used in the paper can be found at https://github.com/Michaelsqj/capria_motion_correction We are currently unable to share subject level data due to data protection issues, although our center is actively working on a solution to this.

## Author Contributions

Qijia Shen: Conceptualization, Data Curation, Methodology, Investigation, Formal Analysis, Writing Original draft, Writing — Review & Editing. Wenchuan Wu: Conceptualization, Methodology, Writing — Original draft, Writing — Review & Editing, Supervision. Mark Chiew: Methodology, Software, Writing — Review & Editing. Yang Ji: Data Curation, Writing — Review & Editing. Joseph G. Woods: Methodology, Writing — Review & Editing, Supervision. Thomas W. Okell: Conceptualization, Methodology, Investigation, Formal Analysis, Writing — Original draft, Writing Review & Editing, Supervision, Project administration, Funding acquisition.

## Acknowledgements

We are grateful for funding support from a Sir Henry Dale Fellowship jointly funded by the Wellcome Trust and the Royal Society (220204/Z/20/Z). The Wellcome Centre for Integrative Neuroimaging is supported by core funding from the Wellcome Trust (203139/Z/16/Z) with additional support from the NIHR Oxford Health Biomedical Research Centre (NIHR203316). W.W. is supported by the Royal Academy of Engineering (RF\201819\18\92). MC is supported by the Canada Research Chairs program. The views expressed are those of the authors and not necessarily those of the NIHR or the Department of Health and Social Care. Many thanks also to Jeff Fessler, Philipp Ehses and colleagues for making available their excellent NUFFT and Siemens raw data reading MATLAB code, as well as to Siemens Healthineers for providing the base pulse sequence code that we built upon in this work. For the purpose of open access, the author has applied a CC BY public copyright license to any Author Accepted Manuscript version arising from this submission.

## Supplementary material for ‘Motion correction with subspace-based self-navigation for combined angiography, perfusion and structural imaging’

**Figure S1.**
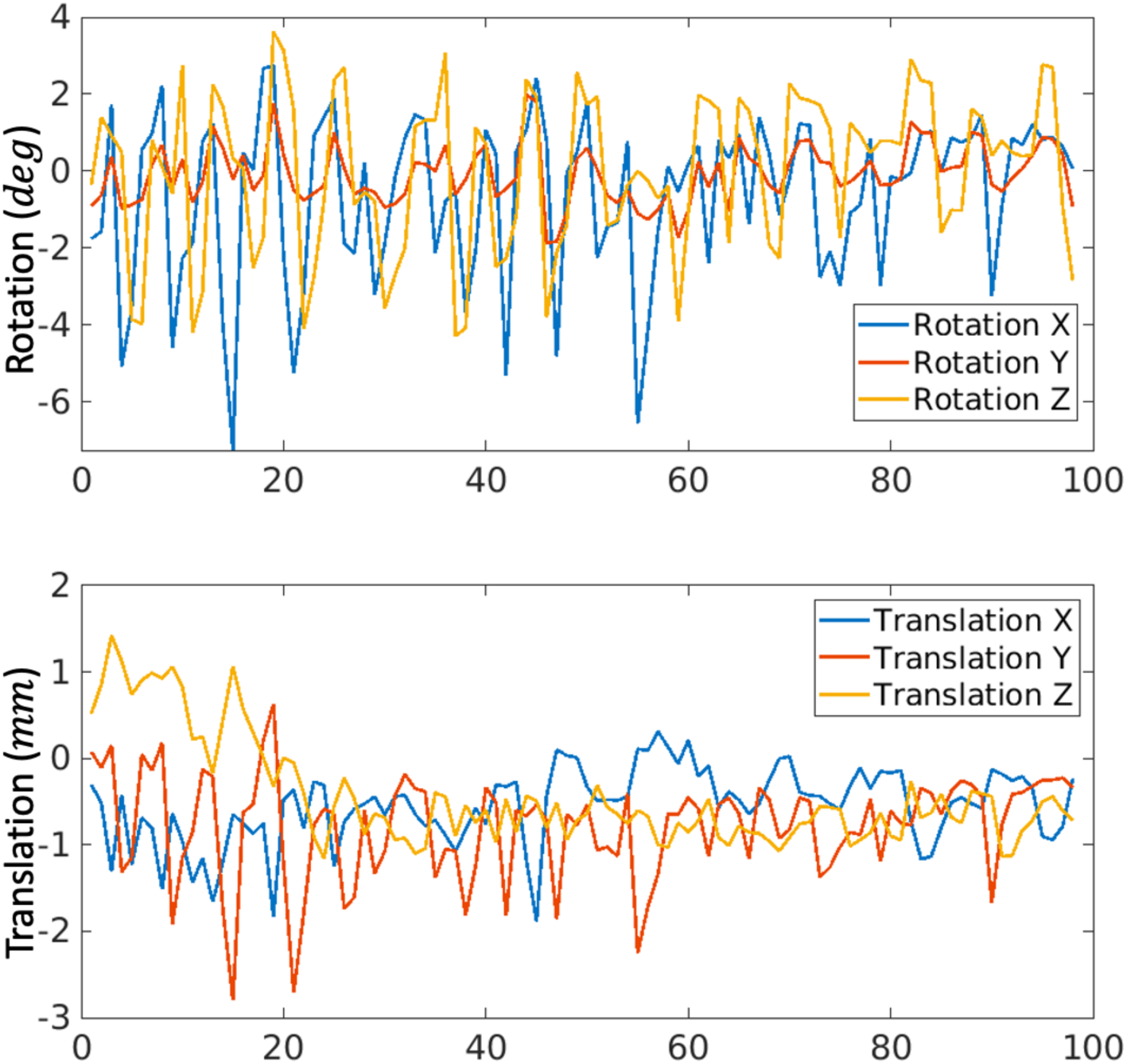
Motion parameters used for numerical simulations.

**Figure S2.**
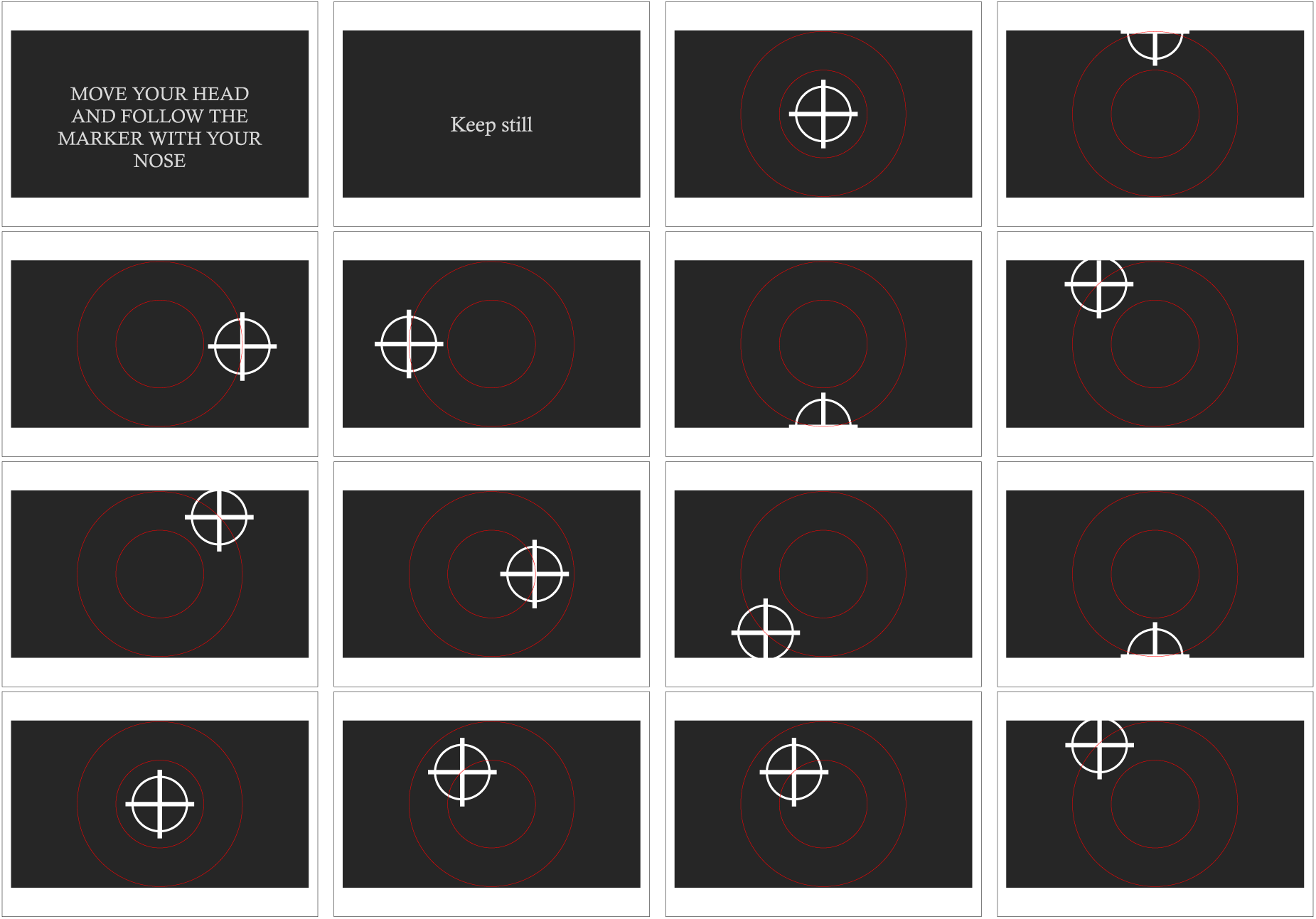
Slides used during cued motion scan. Subjects were instructed to follow the marker on the screen in the scanner.

**Figure S3.**
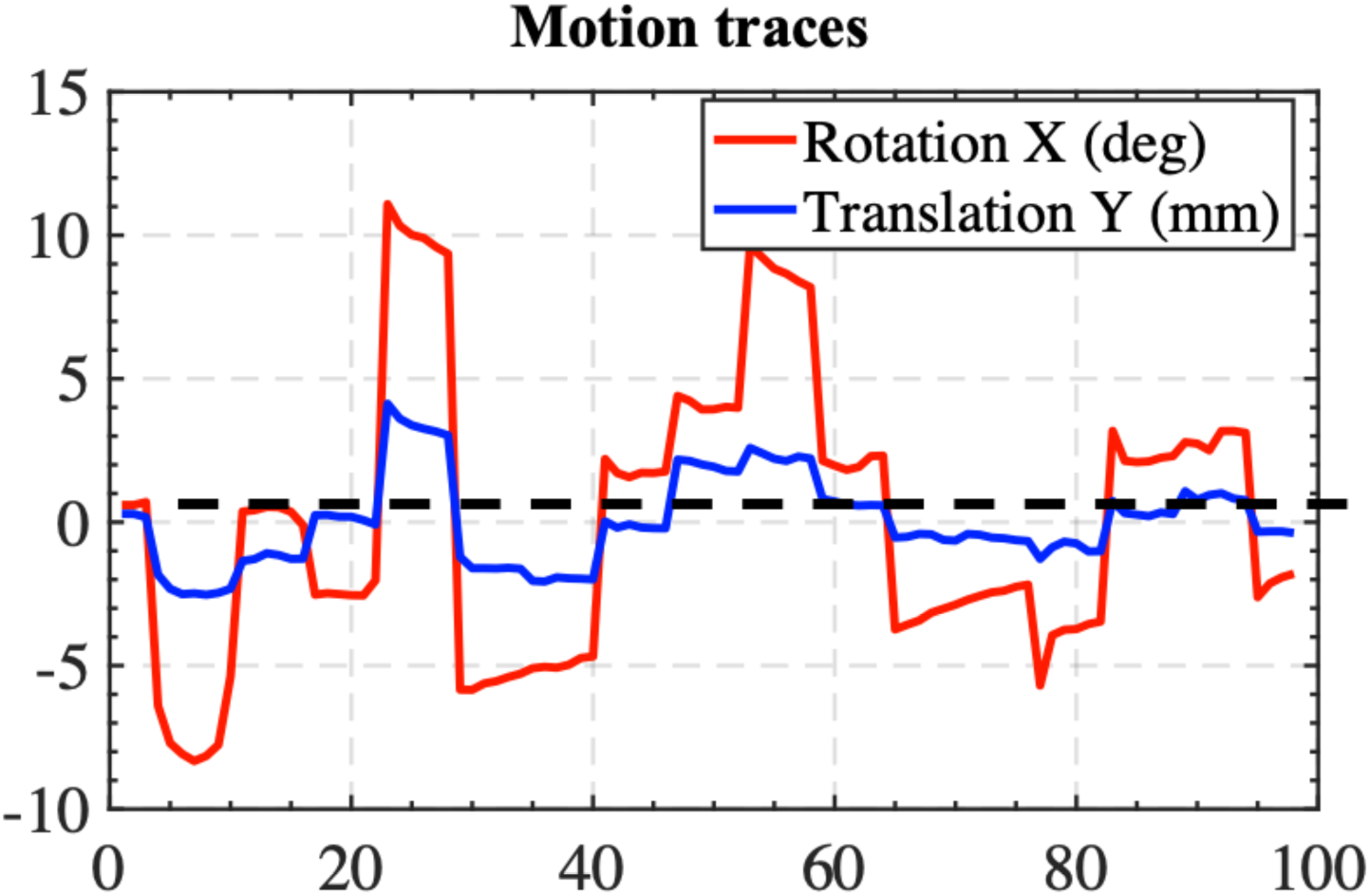
Motion traces for severe subject motion during one acquisition corrupting the angiography images in Figure 5c.

**Figure S4.**
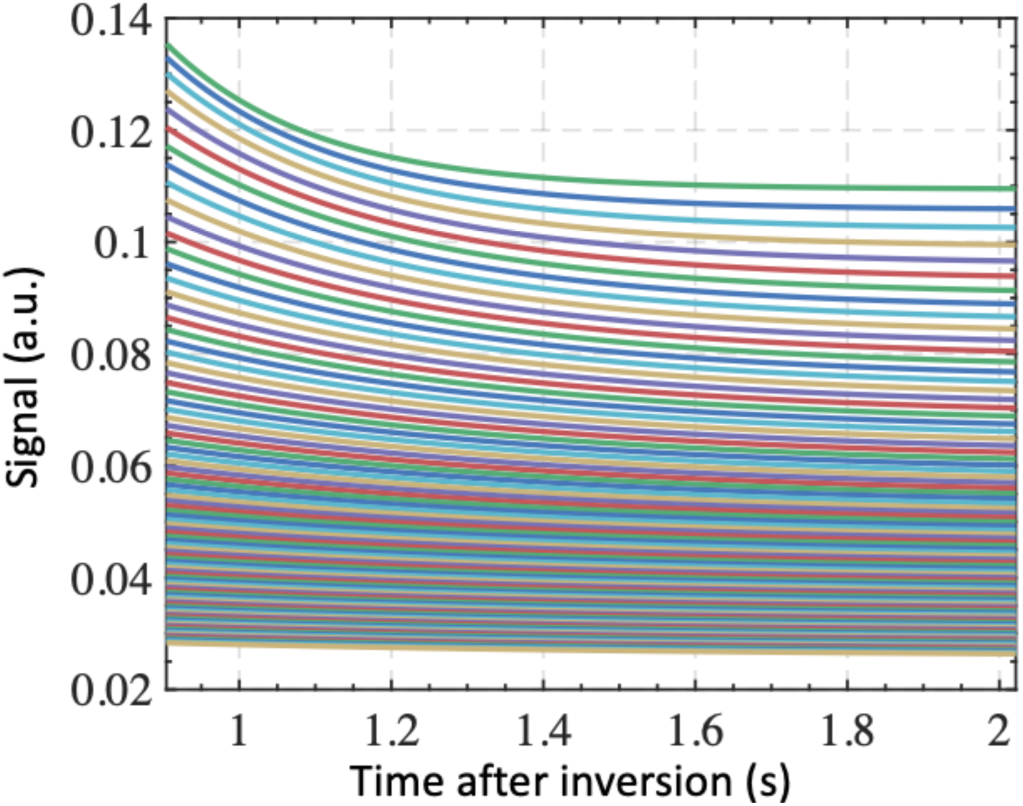
Simulated signal curves for train of readouts in MP-RAGE sequence with TI=904ms, echo train length = 128, echo spacing = 8.8 ms, flip angle = 8°, and T1 ranging from 250∼4000 ms.

## ADMM for joint label and control reconstruction

Here we demonstrate the ADMM algorithm experimented for joint label-control reconstruction with LLR regularization on the label-control difference image.

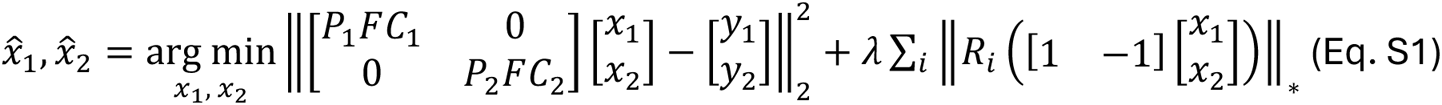

*P*_1_*FC*_1_, *P*_2_*FC*_2_ represents k-space sampling, Fourier transform, and coil sensitivity corresponds to repeats with label and control. *x*_1,_ *x*_2_ represents label and control images to reconstruct. *y*_1_, *y*_2_ represents k-space data of repeats with label and control. *R_i_* represents operation for extracting LLR patches as in Equation 4.

For convenience, Equation S1 is rewritten as

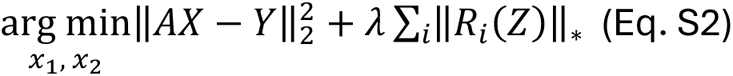

where,

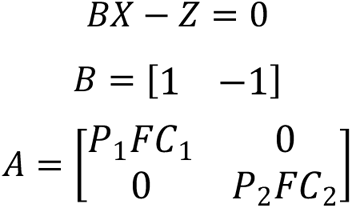

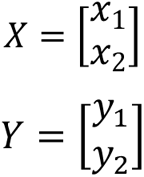

And ADMM updates become

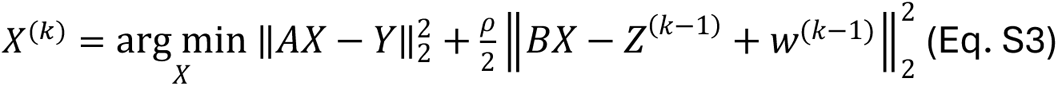

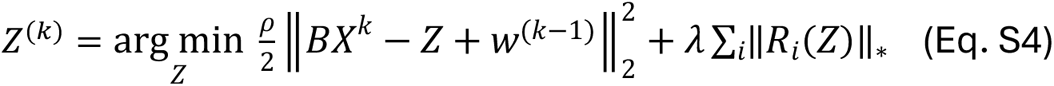

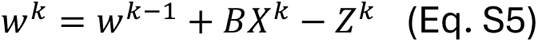

*ρ* and *λ* are two parameters to adjust and *w* is a variable introduced by the Augmented Lagrangian method.

Equation S3 could be solved using the iterative conjugate gradient method (Barrett et al., 1994) which is the most time consuming step in ADMM updates due to the iterative NUFFT (Fessler & Sutton, 2003) calculation, but is prohibited using the method for reconstructing dynamic angiography and perfusion images. Equation S4 is solved by applying locally low rank denoising on *Z*.

